# De novo lipogenesis pathway is a vulnerability in IDH1 mutant glioma

**DOI:** 10.1101/2023.11.15.567231

**Authors:** Lubayna S Elahi, Michael C Condro, Riki Kawaguchi, Yue Qin, Alvaro G. Alvarado, Brandon Gruender, Haocheng Qi, Tie Li, Albert Lai, Maria G. Castro, Pedro R. Lowenstein, Matthew C Garrett, Harley I. Kornblum

**Affiliations:** Department of Psychiatry and Behavioral Sciences and the UCLA Intellectual and Developmental Disabilities Research Center, David Geffen School of Medicine, UCLA, Los Angeles, CA, USA; Department of Molecular and Medical Pharmacology, David Geffen School of Medicine, UCLA, Los Angeles, CA, USA; Department of Neurology, David Geffen School of Medicine, UCLA, Los Angeles, CA, USA; Department of Neurosurgery, Department of Cell and Developmental Biology, and Rogel Cancer Center, University of Michigan Medical School, Ann Arbor, MI, USA; Kettering Health Network Kettering, Ohio, USA

**Author notes:** Corresponding Author: Harley Kornblum, M.D., Ph.D. NRB Room 375D, UCLA 635 Charles E. Young Drive South Los Angeles, CA 90095.

## Abstract

Histone deacetylases (HDACs) have a wide range of targets and can rewire both the chromatin and lipidome of cancer cells. In this study, we show that valproic acid (VPA), a brain penetrant anti-epileptic and histone deacetylase inhibitor, inhibits the growth of IDH1 mutant tumors in vivo and in vitro, with at least some selectivity over IDH1 wild type tumors. Surprisingly, genes upregulated by VPA showed no change in chromatin accessibility at the promoter, but there was a correlation between VPA downregulated genes and diminished promoter chromatin accessibility. VPA inhibited the transcription of lipogenic genes and these lipogenic genes showed significant decrease in promoter chromatin accessibility only in the IDH1 MT glioma cell lines tested. VPA targeted a key lipogenic gene, fatty acid synthase (FASN), via inhibition of the mTOR pathway and both VPA and a selective FASN inhibitor TVB-2640 rewired the lipidome and promoted apoptosis in an IDH1 MT but not in an IDH1 WT glioma cell line. We further find HDACs are involved in the regulation of lipogenic genes and in particular HDAC6 is important for regulation of FASN in IDH1 MT glioma. Finally, we show that FASN knockdown alone and VPA in combination with FASN knockdown significantly improved the survival of mice in a IDH1 MT primary orthotopic xenograft model in vivo. We conclude that targeting fatty acid metabolism through HDAC inhibition and/or FASN inhibition may be a novel therapeutic option in IDH1 mutant gliomas.

## Introduction

Missense mutations in the gene isocitrate dehydrogenase one (IDH1) reprogram the metabolic and epigenetic landscape of IDH1 mutant (MT) gliomas making them molecularly and physiologically distinct from IDH1 wildtype (WT) gliomas. WT IDH1 catalyzes the conversion of isocitrate to alpha-ketoglutarate (α-kg) whereas MT IDH1 reduces α-kg to the oncometabolite 2-hydroxyglutarate (2-HG). 2-HG by blocking α-kg, inhibits the enzymatic function of ten-eleven translocation family of 5-methylcytosine hydroxylases, and Jumanji family of histone demethylases resulting in an increase in both histone and DNA methylation^1,2^. Inhibitors that directly target MT IDH1 and block 2-HG have efficacy in low grade IDH1 mutant gliomas but have not been proven to be beneficial for mutant high-grade gliomas ^3–6^.

Chromatin modifying drugs such as histone deacetylase inhibitors (HDACis) are promising candidates for therapeutic approaches for gliomas and have been shown to inhibit growth of glioma by multiple mechanisms such as cell cycle arrest, apoptosis, demethylation, and metabolic rewiring^7–9^. Broad spectrum HDACis target histones and alter the chromatin but can also target non-histone proteins which in an HDAC dependent or independent manner can alter cellular signaling and metabolism^7^. In recent years, one non histone protein that has emerged as an interesting target of HDACis is fatty acid synthase (FASN). Acetylation destabilizes FASN protein and results in inhibition of de novo lipogenesis^10^. FASN is a key enzyme in the de novo lipogenesis pathway and catalyzes the synthesis of palmitate from acetyl-coA, malonyl-coA and nicotinamide adenine dinucleotide phosphate (NADPH). High 2-HG in IDH1 MT tumors increases the consumption of NADPH and limits the availability of NADPH for de novo lipogenesis^11,12^, raising the possibility that such tumor cells could be selectively vulnerable to FASN inhibition in IDH1 MT tumors.

In this study, we show that IDH1 MT gliomas are sensitive to VPA treatment. We characterized the effect of VPA on the overall chromatin architecture of IDH1 MT gliomas and show that VPA alters promoter accessibility and inhibits transcription of several lipogenic enzymes including FASN in IDH1 MT gliomas. We show both VPA and the selective FASN inhibitor TVB-2640 rewires the lipidome and promotes apoptosis in a IDH1 MT but not IDH1 WT glioma cell line. Finally, since cancer cells can easily rewire their metabolism to support growth, we propose targeting fatty acid metabolism by HDAC and FASNi alone or in combination is a novel therapeutic option for IDH1 MT gliomas.

## Results

### IDH1 MT glioma are sensitive to VPA in vitro and in vivo

We tested three different HDACis VPA, panobinostat, and belinostat in varying concentrations in IDH1 WT and IDH1 MT primary glioma cell lines. We found IDH1 MT glioma cell lines had a slightly lower IC50 for VPA compared to IDH1 WT cell lines (WTIC50=1.23; MTIC50=0.96) and the hill slope between the two dose response curves were statistically significant (WTHill Slope =-0.80; MTHill Slope=-1.68; 0.0032) (Fig1A). Panobinostat (Fig1B) inhibited growth of both IDH1 WT and IDH1 MT cell lines but there was no significant difference in the IC50 or hill slope between the two dose response curves. The IDH1 WT cell lines tested were more sensitive to belinostat than IDH1 MT glioma cell lines (WTIC50=0.84; MTIC50=1.97) (Fig1C). As expected the heterozygous IDH1 MT cell lines showed significantly higher 2HG compared to a IDH1 WT cell line (Fig1D). The 2HG content in hemizygous IDH1 MT cell line BT142 was approximately double that of the IDH1 WT cell line but the difference did not reach statistical significance. The heterozygous IDH1 MT cell lines HK252 and HK211 do not form xenograft in vivo even in NSG mice; hence we tested the efficacy of VPA in vivo in the hemizygous BT142 model. Although BT142 has limited 2HG production (Fig1D), we found that the transcriptome of BT142 is most similar to our heterozygous IDH1 MT cell line HK252 and still distinct from IDH1 WT cell lines (SFig1A), indicating that even with the loss of 2HG production, the line possesses many of the essential features of IDH1 mutant gliomas. We found in vivo VPA treatment significantly improved survival of mice in the BT142 xenograft model by 17 days (Fig1E).

**Fig1:**
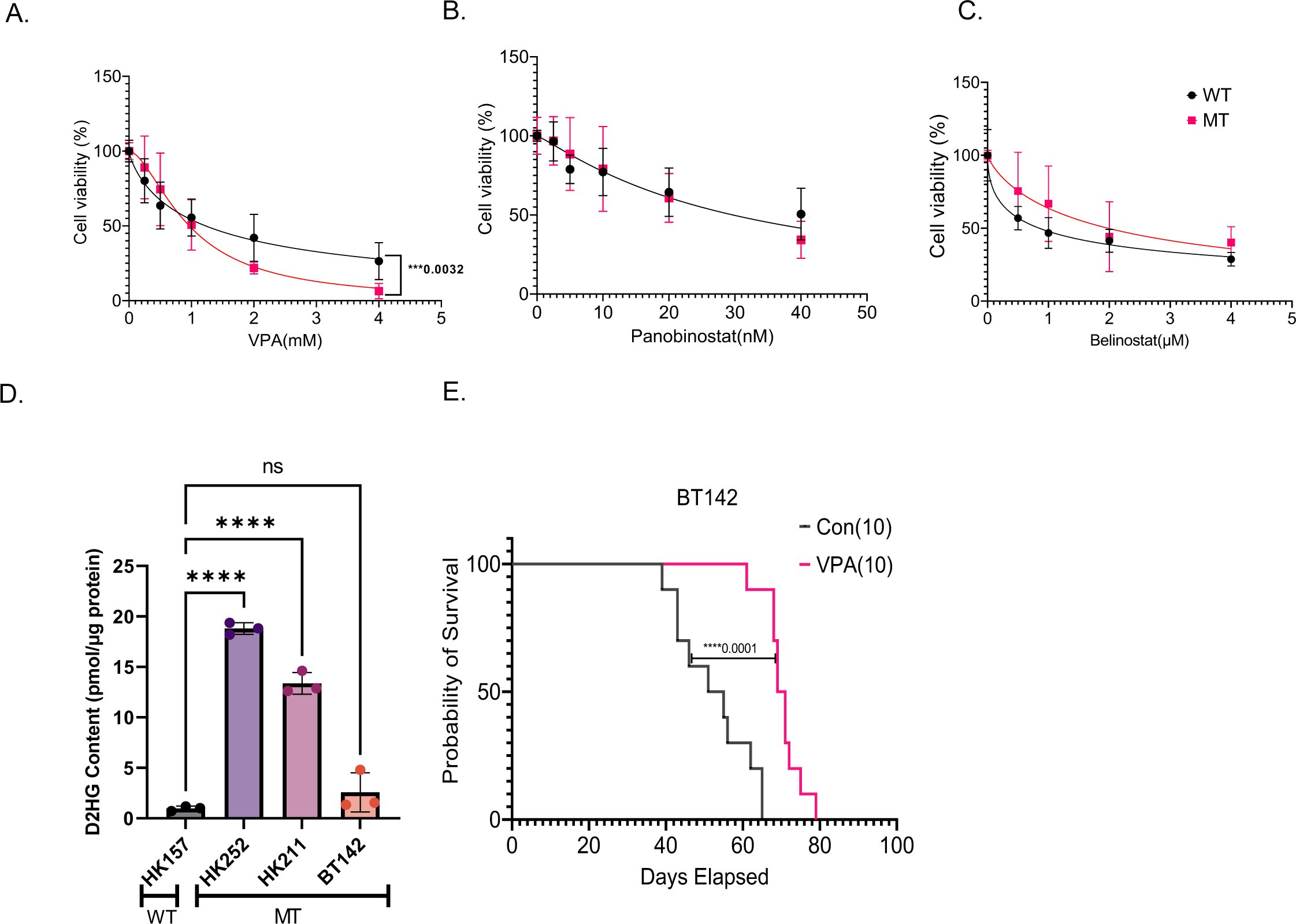
IDH MT GBM are sensitive to VPA in vitro and in vivo. A-C Dose response curve of IDH MT (3 cell lines) and IDH WT(2 cell lines) treated with VPA (A), Panobinostat (B), and Belinostat (C) for 1 week. D. D-2HG content in the human primary WT & MT cell lines. E. Kaplan-Meier survival curve for mice treated with either saline or VPA (300mg/kg, twice daily) implanted with BT142. Non-Linear Regression, One way ANOVA,****P value<0.0001; ***P value<0.001; Error bars ±SD

Next, we tested the efficacy of HDACis in a syngeneic murine IDH1 (mIDH1) glioma model. In vitro, NPAIC1(NRAS/IDH11^R132H^/shP53/shATRX) compared to mIDH1 WT cell line NPAC54B (NRAS/shP53/shATRX) had a modestly lower IC50 for both VPA (NPAC54BIC50=1.65; NPAIC1IC50=1.37) (Fig2A) and belinostat (NPAC54BIC50=0.57; NPAIC1IC50=0.28) (Fig2B). The mIDH1 MT cell line NPAIC1 also had significantly higher 2HG than mIDH1 WT cell line NPAC54B (Fig 2C).

**Fig2.**
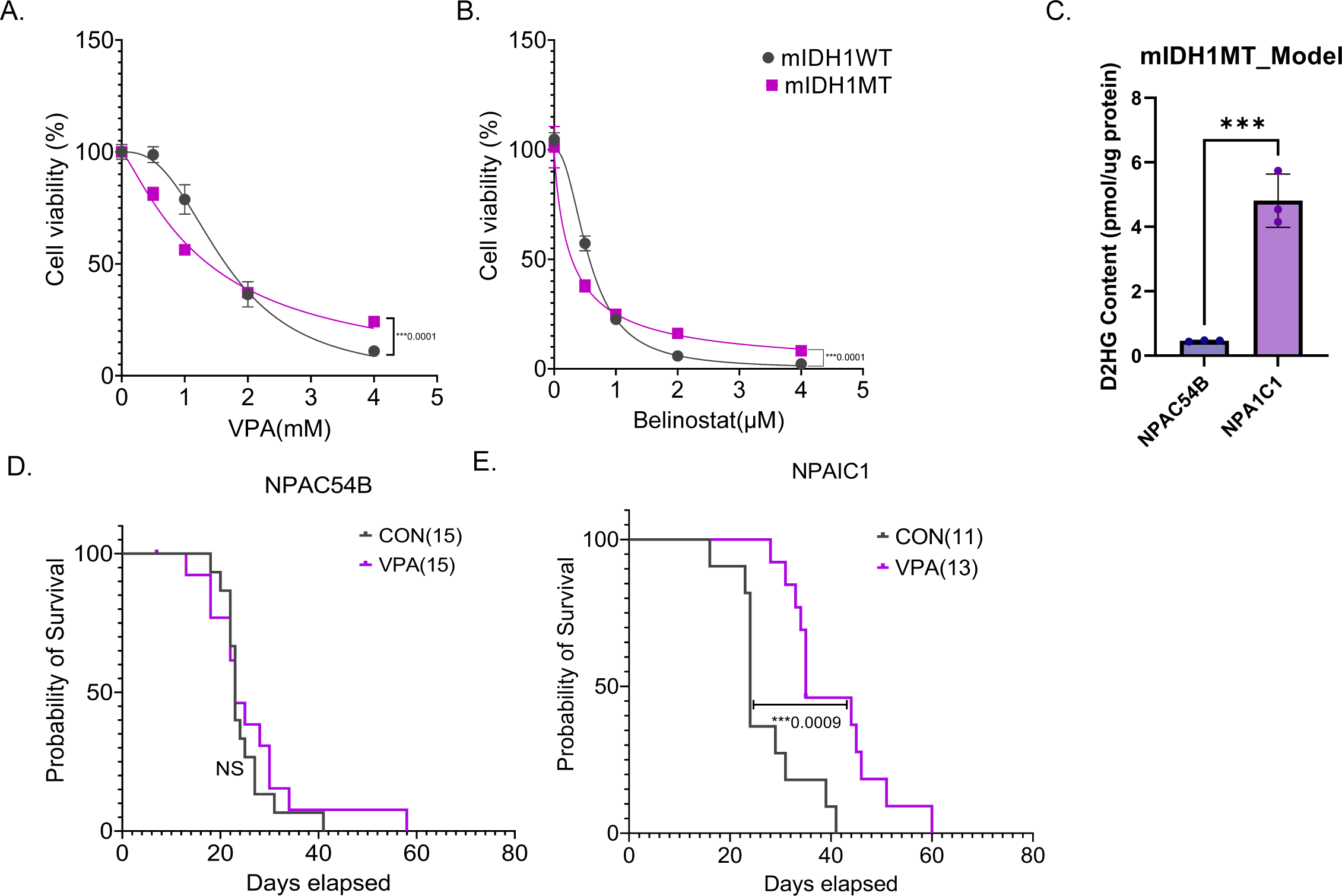
VPA improves survival of mice in a murine isogenic model of IDH MT GBM. A-B Dose response curve of mIDH1MT and mIDH1WT treated with VPA (D) or Belinostat(E) for 5days. C. D-2HG content in murine mIDH1WT and mIDH1MT cell lines NPAC54B and NPAIC1. D-E. Kaplan-Meier survival curve for mice with NPAC54B or NPAIC1 xenograft treated with either saline or VPA (300mg/kg, twice daily). F-G Kaplan-Meier survival curve for mice with NPAC54B or NPAIC1 xenograft treated with either saline, VPA (300mg/kg, twice daily), AG881(2mg/kg, twice daily) or both. Non-Linear Regression, T-test, ****P value<0.0001; ***P value<0.001, **P value<0.01; *P value<0.05; Error bars ±SD

In vivo, VPA treatment did not improve survival of mice bearing NPAC54B tumor xenografts (Fig2D) but significantly improved survival of mice bearing NPAIC1 tumor xenografts by 10 days (Fig2E). Thus, in the tumor models that we examined, we established that VPA diminished tumor growth in vivo and in vitro, but the murine model suggests a greater efficacy in IDH1 mutant tumors.

### VPA has opposing effects on overall gene expression and chromatin accessibility

VPA is a multifaceted drug that has several direct^13^ and indirect targets^14^. Given the pleiotropic effects of VPA, we wanted to understand why IDH1 MT glioma cell lines showed some selectivity to VPA treatment. Given that HDACs are a known target of VPA, we treated a IDH1 MT cell line HK252 and a IDH1 WT cell line HK157 with 1mM VPA and examined histone 3 lysine 27 acetylation(H3K27ac) over time. We found that after 96hr of treatment VPA induced a strong increase in H3K27ac in both IDH1 WT (Fig3A) and IDH1 MT (Fig3B) cell lines as would be expected with HDAC inhibition. Semi quantitative WB analysis showed that, compared to control, HK252 had a 7.6-fold increase in H3K27ac after 96hr whereas HK157 only had a 3.1-fold increase in H3K27ac compared to control. We next treated multiple IDH1 MT glioma cell lines (4) and IDH1 WT glioma cell lines (5) with 1mM VPA for 48 hours and conducted bulk RNA sequencing to get a better understanding of the transcriptomic effect of VPA. A greater number of genes were upregulated than downregulated with VPA treatment in both IDH1 WT and IDH1 MT glioma cell lines (Fig3C). To better understand what biological processes are altered by VPA we performed gene set enrichment analysis (GSEA) and compared the gene ontology (GO) terms that were up and downregulated with VPA treatment in IDH1 WT and IDH1 MT cell lines. We found in both IDH1 WT and IDH1 MT primary glioma cell lines GO terms related to cell cycle and DNA repair were downregulated whereas GO terms related to neurogenesis were upregulated after VPA treatment (Fig3D). We next asked what biological processes were only altered in IDH1 MT and not in IDH1 WT after VPA treatment. We found that VPA upregulated biological processes related to differentiation and downregulated lipid and sterol biosynthetic processes in the IDH1 MT but not IDH1 WT cell lines (Fig3E). We examined a panel of lipogenic genes and found that SREBF1, ACLY, FASN, SCD, and HMGCR were significantly downregulated in primary IDH1 MT cell lines after VPA treatment (Fig3F). In the IDH1 MT cell lines VPA inhibited expression of FASN and HMGCR much more than in the IDH1 WT cell lines. We also treated mIDH11WT and mIDH11MT with 1mM VPA for 48hr and conducted RNA sequencing. We found that VPA downregulated FASN in the mIDH11MT but not in the mIDH11WT cell line (SFig2A).

**Fig3:**
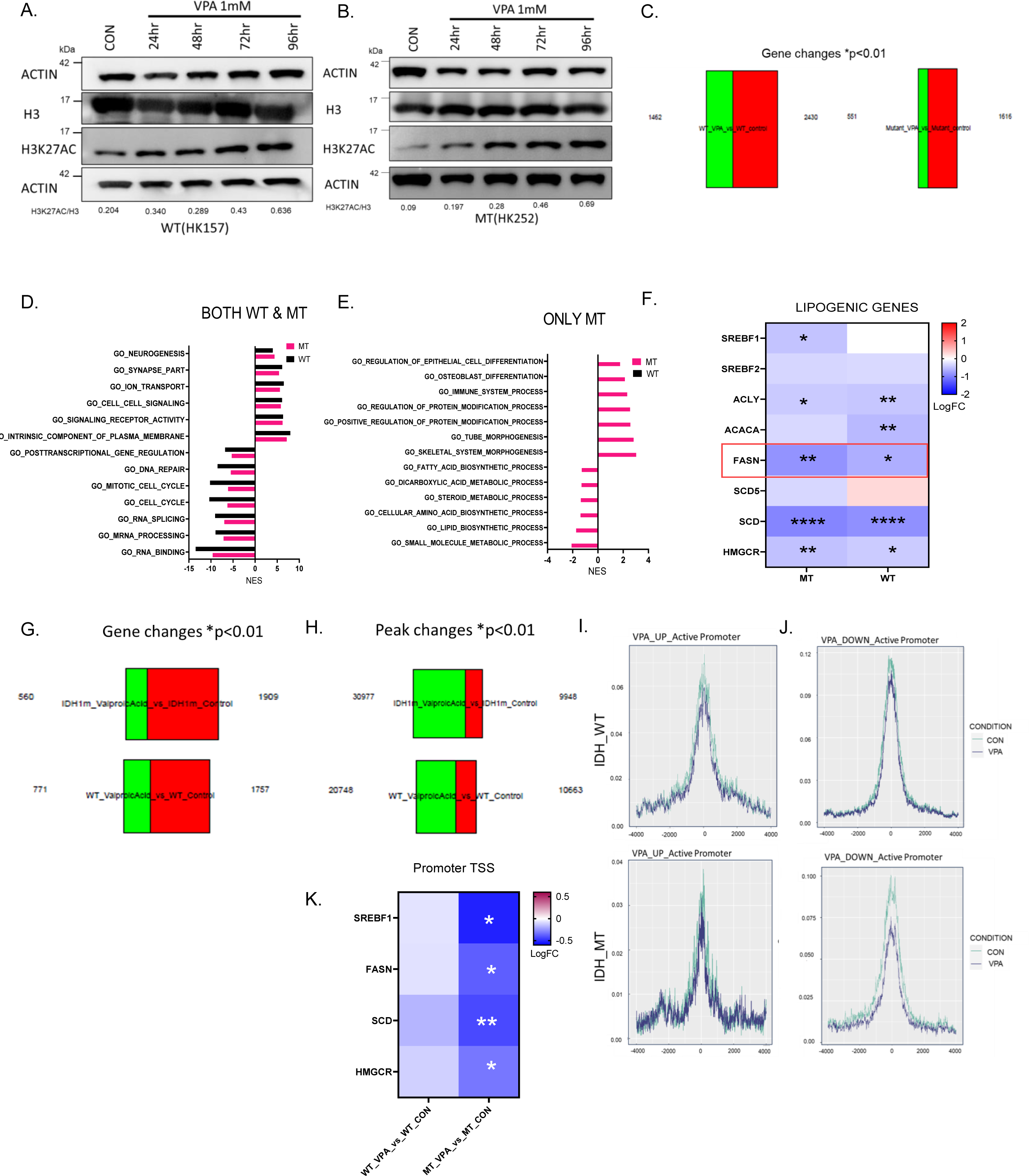
VPA decreases promoter chromatin accessibility and inhibits gene expression of lipogenic genes in IDH MT GBM. A-B. Representative western blot of a IDH WT cell line HK157 (A) and IDH MT cell line HK252 (B) showing increase in H3K27ac over time with treatment of 1mM VPA. C. Differential gene expression changes, red (upregulated genes), green (downregulated genes) in IDH WT (5 cell lines) and IDH MT (4 cell lines) treated with 1mM VPA for 48hours. D-E.. GSEA analysis showing top gene ontology terms upregulated and downregulated in both primary WT and MT cells lines (D) and only in MT but not WT (E). F. Selected lipogenic genes that are downregulated in IDH WT and MT cell lines. G. Differential gene expression changes, red (upregulated genes), green (downregulated genes) in IDH WT (2 cell lines) and IDH MT (3 cell lines) treated with 1mM VPA for 5 days. H. Differential peak changes, red (open peaks), green (closed peaks) in IDH WT (2 cell lines) and IDH MT (3 cell lines) treated with 1mM VPA for 5 days. I-J Active peak changes at promoter (TSS=0) of VPA upregulated and downregulated genes in IDH WT (I) and IDH MT (J) GBM cell lines. K. Selected lipogenic genes that shows alteration in promoter chromatin accessibility in the MT but not WT cell lines. ****P value<0.0001; ***P value<0.001; **P value<0.01; *P value<0.05.

We next re-analyzed our previously published RNA-seq and ATAC-seq dataset^15^ of IDH1 MT and IDH1 WT cell lines treated with 1mM VPA for 5 days and found that similar to the current findings, VPA treatment increased a greater number of genes and downregulated a smaller number of genes in both IDH1 WT and IDH1 MT cell lines (Fig3G). Interestingly, ATAC seq analysis showed that despite of the relatively increased transcription, a greater number of peaks were lost and only a few peaks were gained after VPA treatment (Fig3H), suggesting VPA treatment activates transcription but paradoxically results in chromatin condensation. We next integrated ATAC and RNA seq data and when we examined the promoter region of VPA-upregulated genes in both IDH1 WT and IDH1 MT cell lines, we found that most genes did not show change in promoter accessibility (Fig3I). There was a stronger correlation between promoter chromatin accessibility and VPA downregulated genes in IDH1 MT compared to the IDH1 WT cell lines (Fig3J). Interestingly, promoter accessibility of lipogenic genes were only significantly altered in the IDH1 MT but not IDH1 WT cell lines (3K). Taken together, our findings thus far indicate that VPA, while inhibiting the growth of both IDH1 mutant and wildtype cells has differential molecular effects on the IDH1 mutant and wildtype cells that we examined.

### mTOR pathway is involved in downregulation of FASN by VPA

mTOR signaling plays an important role in regulating lipogenesis^16^ and VPA has been found to inhibit the mTOR pathway in various cancer cell lines^17,18^ so we wondered whether VPA may be inhibiting lipogenic genes via the mTOR pathway. FASN is a key enzyme in the de novo lipogenesis pathway and although VPA targeted several enzymes involved in lipid metabolism VPA inhibited transcription of FASN significantly more in the IDH1 MT compared to IDH1 WT. In addition, both in our IDH1 MT primary tumor cell lines and murine model, VPA inhibited transcription of FASN; hence we focused our attention on FASN.

We treated IDH1 WT cell line HK157 and 2 IDH1 MT glioma cell lines, HK252 and BT142 with VPA for four days and examined phosphorylation of ribosomal protein S6 (PS6) and FASN protein expression. We found that VPA treatment had a minimal effect on PS6 and FASN protein expression in the IDH1 WT cell line HK157 (Fig4A) but resulted in diminished PS6 and FASN protein expression in IDH1 MT cell lines HK252 (Fig4B) and BT142 (Fig4C). Treatment of MT cell line HK252 with rapamycin (RAPA) also diminished PS6 and FASN protein expression (Fig4D). In HK252, the combination of VPA with RAPA was better than VPA alone in decreasing FASN protein expression (Fig4E). RAPA treatment also inhibited FASN mRNA expression in HK252 while the addition of VPA to RAPA did not further diminish mRNA expression of FASN in HK252 (Fig4F). Treatment with RAPA inhibited growth of IDH1 MT cell lines (Fig4G). Interestingly, the combination of VPA with RAPA was not better than VPA alone in inhibiting growth in HK252, but in HK157 and BT142 the combination of VPA and RAPA was better than VPA alone in inhibiting growth (Fig4G) suggesting there is some heterogeneity in terms of how cell lines respond to mTOR inhibition. We also tested the effect of VPA and RAPA on NPAC54Band NPAIC1. Surprisingly, VPA increased PS6 in NPAC54B but decreased PS6 in the mutant NPAIC1 cell line in a dose dependent manner (SFig3A). Treatment with RAPA inhibited the growth of only NPAIC1 but not NPAC54B. The combination of VPA and RAPA was better at inhibiting growth of NPAC54B but not NPAIC1 cell line (SFig3B). In summary, we conclude that in both the IDH1 MT human primary tumor cell lines and in the murine model VPA inhibits growth at least partially via inhibition of the mTOR pathway and that the mTOR pathway is involved in downregulation of FASN. However, there is line to line heterogeneity in the relative extent to which FASN and mTOR signaling are connected.

**Fig4:**
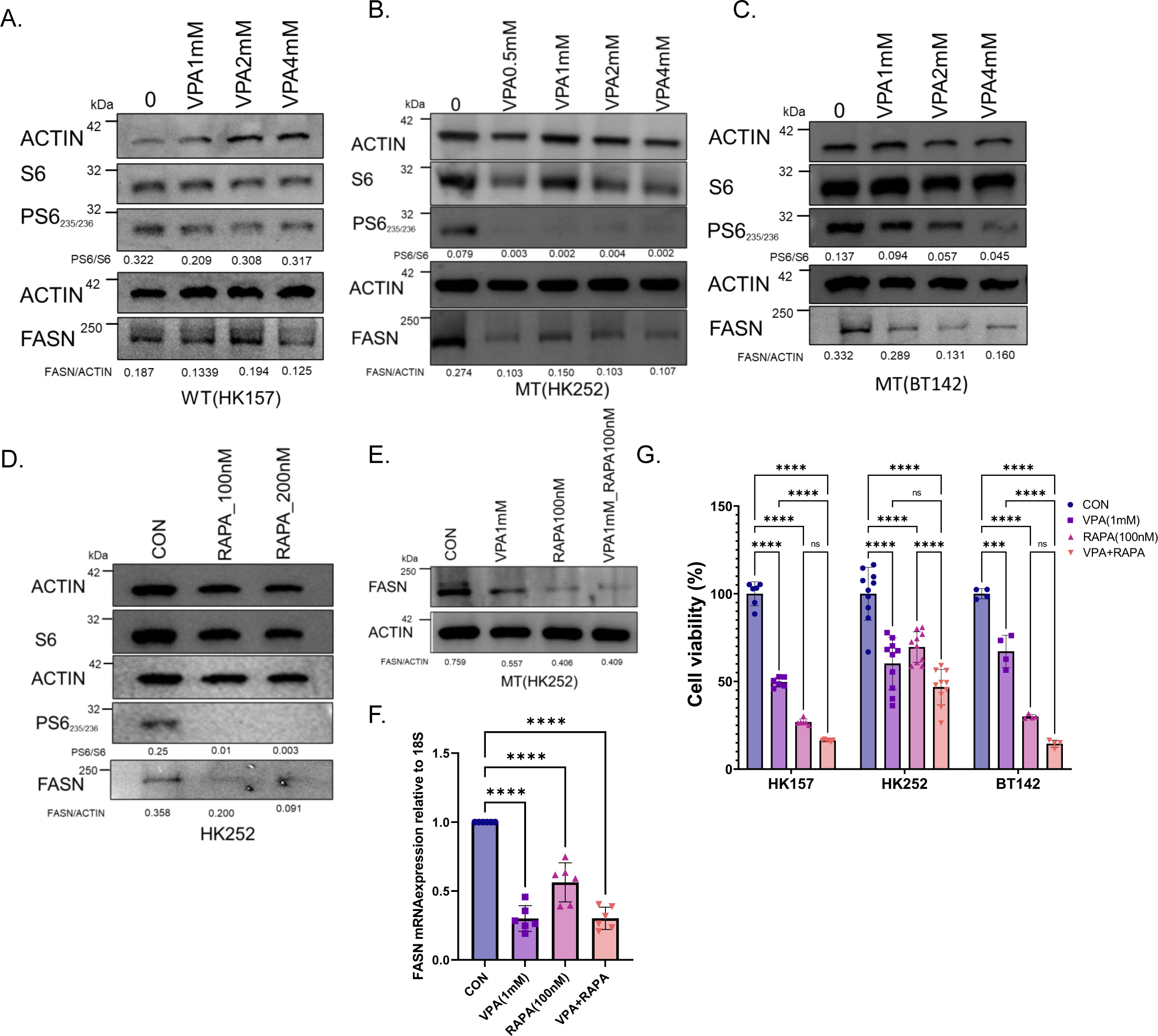
VPA may regulate FASN mRNA and protein through inhibition of mTOR signaling. A-C. Representative western blots of PS6 and FASN in a IDHWT GBM cell line HK157 (A), and IDH MT GBM cell lines HK252 (B) and BT142 (C) treated with different doses of VPA for 4 days. D. Representative western blot of PS6 and FASN in MT cell line HK252 treated with rapamycin for 4 days. E. Representative western blot of PS6 and FASN in MT cell line HK252 treated with VPA, rapamycin, & the combination for 4 days. F. Relative mRNA expression of FASN in MT cell line HK252 treated with VPA, rapamycin, & the combination for 4 days. H. Relative cell viability after treatment with VPA, RAPA, & combination for 1 week. 2-way ANOVA and post hoc t-test ****P value<0.0001; ***P value<0.001; Error bars ±SD

### Inhibition of FASN inhibits the growth of IDH1 mutant cell lines

Next to test whether direct inhibition of FASN has any effect on growth of IDH1 MT cell lines we treated IDH1 WT and MT cell lines with a selective FASN inhibitor (FASNi) TVB-2640 (Fig5A). We found that TVB-2640 inhibited the growth of WT cell line HK157(Fig5B) and MT cell lines HK252(Fig5C) and BT142(Fig5D). Supplementation of palmitate in the media rescued the growth inhibitory effect of TVB-2640 in both WT and MT cell lines, indicating that the negative effects on growth were selectively mediated by FASN inhibition. In the murine model, TVB-2640 inhibited the growth of both NPAC54B and NPA1C1 (SFig4A). Thus, while FASN expression is more sensitive to VPA treatment in IDH1 mutant cells, both IDH1 MT and IDH1 WT cells are sensitive to direct FASN inhibition.

**Fig5:**
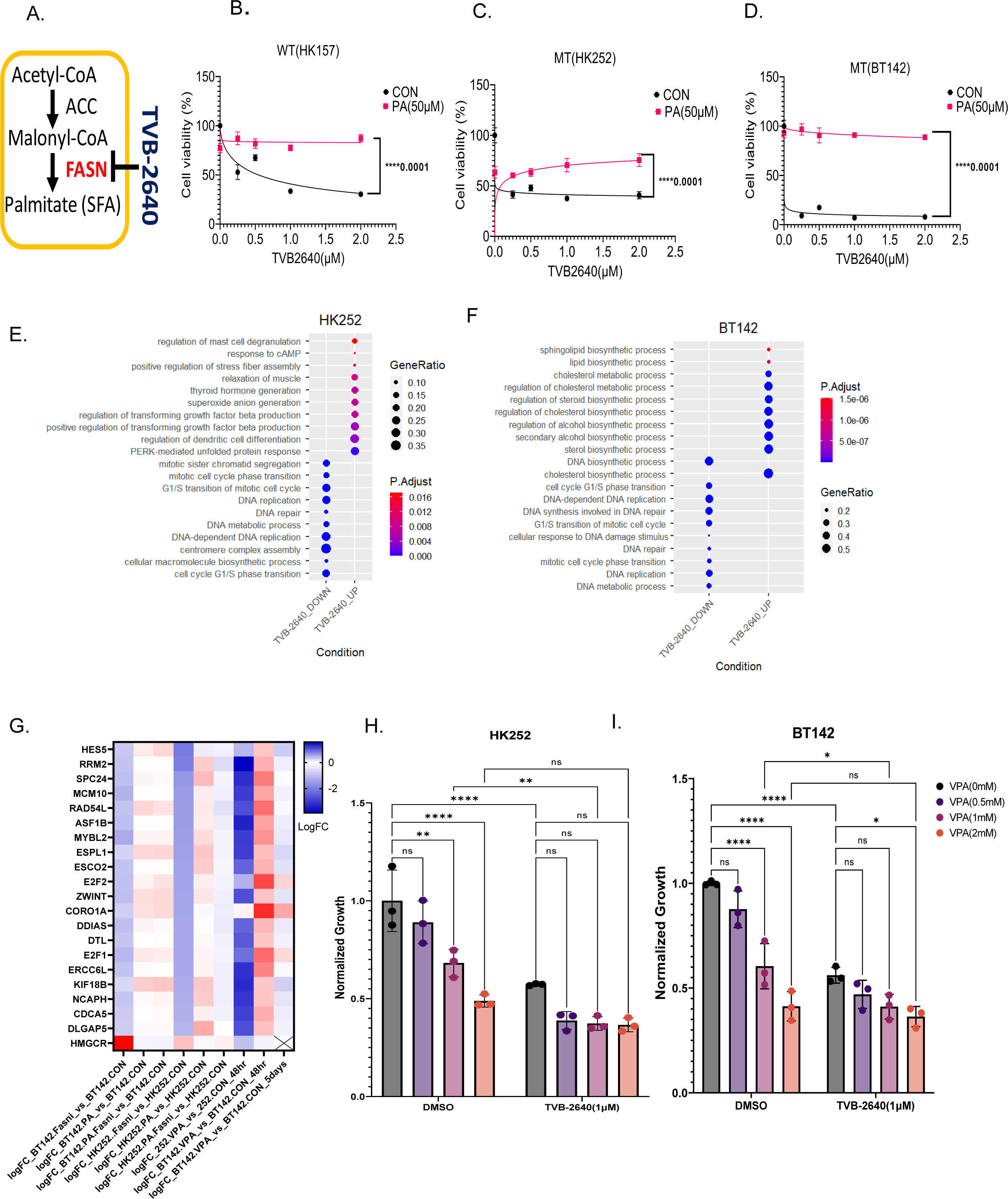
Selective FASN inhibitor TVB-2640 inhibits cell cycle genes in MT cell lines which can be recovered by supplementation of palmitate. A. Schematic of TVB-2640 inhibiting FASN. B-D. Relative cell viability of WT cell line HK157(B) and MT cell lines HK252(C), BT142(D) treated with TVB-2640 in the presence and absence of palmitate for 1 week. Non-Linear Regression, ****P value<0.0001; E-F. Dot plot showing upregulated and downregulated GO terms for MT cell lines HK252 (B) and BT142(C) after 4 days of treatment with TVB-2640(1µM). G. Heat map of common genes inhibited by TVB-2640 and VPA in both HK252 and BT142. H-I Relative growth of HK252 and BT142 treated with VPA (1mM), TVB-2640 (1µM), and PA (50µM) for 1 week. 2-way ANOVA, post hoc t-test, ****P value<0.0001; ***P value<0.001; **P value<0.01; **P value<0.05; Error bars ±SD

To better understand the effects of FASN inhibition, we treated IDH1 MT cell lines with TVB-2640, palmitate, and the combination for 4 days and performed RNA sequencing. We found that TVB-2640 downregulated biological processes related to DNA repair and cell cycle in both HK252 (Fig5E) and BT142(Fig5F). Genes related to cell cycle such as (E2F1, E2F2, RRM2) and anti-apoptosis (DDIAS, MCM10) were downregulated by TVB-2640 and the expression of these genes could be recovered by palmitate supplementation (Fig5G). However, we noticed that TVB-2640 upregulated different biological processes in HK252 and BT142. In HK252 biological processes such as TGFβ production, stress fiber assembly and unfolded protein response were upregulated whereas in BT142 sphingolipid and cholesterol biosynthetic process were upregulated after FASN inhibition. When we looked at individual genes, we found 3-hydroxy-3-methylglutaryl-CoA reductase (HMGCR) was upregulated in both HK252 and BT142. Treatment with lovastatin, an HMGCR inhibitor inhibited growth of both MT cell lines HK252 and BT142 (SFig5A). The combination of lovastatin with TVB-2640 was better at inhibiting growth of HK252 (SFig5B). This suggests IDH1 MT GBM cell lines perhaps increase cholesterol biosynthesis to compensate for FASN inhibition. When we compared genes that were targeted by both TVB-2640 and VPA we found a suite of genes such as E2F1, E2F2, RRM2, DDIAS, MCM10 were downregulated by both drugs. In BT142 these genes decreased after 5 days of treatment. However, VPA inhibited expression of HMGCR while TVB-2640 upregulated HMGCR in IDH1 MT cell lines. Together we conclude in IDH1 MT glioma cell lines, VPA may work through FASN to inhibit cell cycle and anti-apoptotic genes. However, while VPA treatment inhibited transcription of several lipogenic genes, TVB-2640 treatment did not inhibit all lipogenic genes but upregulated some lipogenic genes such HMGCR and ACACA suggesting the metabolic changes induced by VPA and TVB-2640 may be different.

Lastly, we wanted to assess the combinatorial effect of TVB-2640 and VPA. We treated HK252 (Fig5H) and BT142 (Fig5I) with VPA and TVB-2640 and the combination of both drugs for 1 week. In HK252, the combination of VPA with TVB-2640 was not significantly better than TVB-2640 (1µM) alone in inhibiting growth. In BT142, the combination of VPA2mM with TVB-2640 (1µM) but not the combination of VPA(1mM) with TVB-2640(1µM) was significantly better than TVB-2640 (1µM) alone in inhibiting growth. Although we don’t see a rescue the lack of additivity of the two drugs suggests at least part of the effect of VPA is mediated via FASN.

### VPA and TVB-2640 have distinct effect on free fatty acids in IDH1 MT GBM cell line

The primary product of FASN is palmitate. Therefore, we next conducted lipidomics to measure palmitate and other free fatty acids after VPA and TVB-2640 treatment. In the IDH1 WT cell line, HK157 treatment with both VPA (Fig6A) and TVB-2640 (Fig6B) for 4 days significantly decreased free palmitic and stearic acid. Surprisingly, treatment with TVB-2640 significantly increased free palmitic and stearic acid in HK252 (Fig6C) suggesting involvement of a compensatory mechanism or mechanisms. VPA treatment, on the other hand, significantly increased oleic acid but did not alter palmitic or stearic acid in IDH1 MT GBM cell line HK252(Fig6D). This further suggests that while VPA targets FASN, the net metabolic effect of VPA and TVB-2640 on the lipidome is distinct in the IDH1 MT GBM cell line and it is different when compared to IDH1 WT cell line HK157. In IDH1 WT cell line HK157, both VPA and TVB-2640 decreased free saturated fatty acids while in HK252 TVB-2640 significantly increased free saturated fatty acids. However, in HK252 VPA shifted the ratio of free fatty acids towards monounsaturated fatty acids suggesting VPA may not solely working via inhibition of FASN.

**Fig6:**
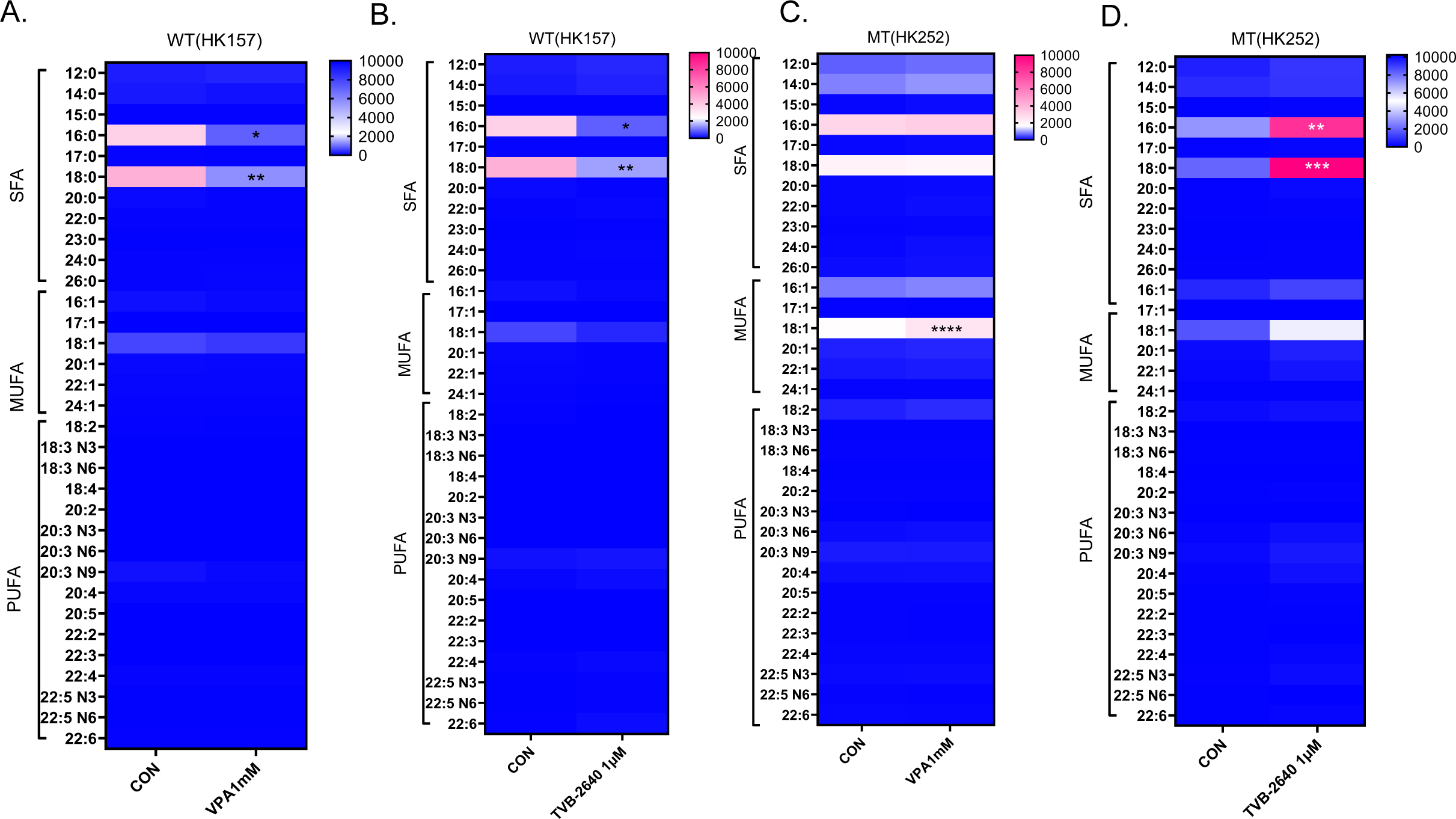
VPA and TVB-2640 increases free fatty acid in a IDH MT GBM but not a IDH WT cell line. A-D. Heatmap showing change in saturated and unsaturated free fatty acids after 4 days of treatment with VPA and TVB-2640; 2-way ANOVA, post hoc t-test, ****P value<0.0001; ***P value<0.001; **P value<0.01; *P value<0.05.

### VPA and TVB-2640 alter lipid droplets and induces apoptosis

To prevent lipotoxicity, excess free fatty acids are often converted into neutral lipids and stored in organelles called lipid droplets. Both oleic acid^19^ and VPA^20^ have been previously found to induce lipid droplets. Our lipidomic data thus far suggested that both VPA and TVB-2640 can increase free fatty acids in IDH1 MT, so we wondered if this might also have an effect on lipid droplet formation. We found in the MT cell line HK252, treatment with VPA for 4 days increased the amount of lipid droplets in lipid droplet positive cells as shown in Fig7B and quantified in Fig7I when compared to control (Fig7A, 7I). There was no difference in lipid droplet formation with TVB-2640 (Fig7C, 7I) but the combination of VPA and TVB-2640 inhibited lipid droplet accumulation by VPA (Fig7D, 7I). Treatment of IDH1 MT cell line HK252 with 100µM oleic acid also increased lipid droplets (Fig7E,7K) and TVB-2640 in combination with oleic acid (Fig7F,7K) inhibited lipid droplet formation. The combination of VPA and oleic acid also increased lipid droplets compared to control (Fig7K). Lipid droplets in the VPA with oleic acid condition appeared much larger and appeared to fuse with each other (Fig7G). Interestingly, unlike HK252, where we found a significant increase in lipid droplets with both VPA (Fig7I) and oleic acid (Fig7K), in HK157 only oleic acid significantly increased lipid droplets (Fig7J) but not VPA(Fig7L). These findings indicate that the increase in lipid droplets in HK252 may be connected to the increase in free oleic acid induced by VPA. However, in HK157 we saw a decrease in free fatty acids, and hence we saw no change in lipid droplets.

**Fig7:**
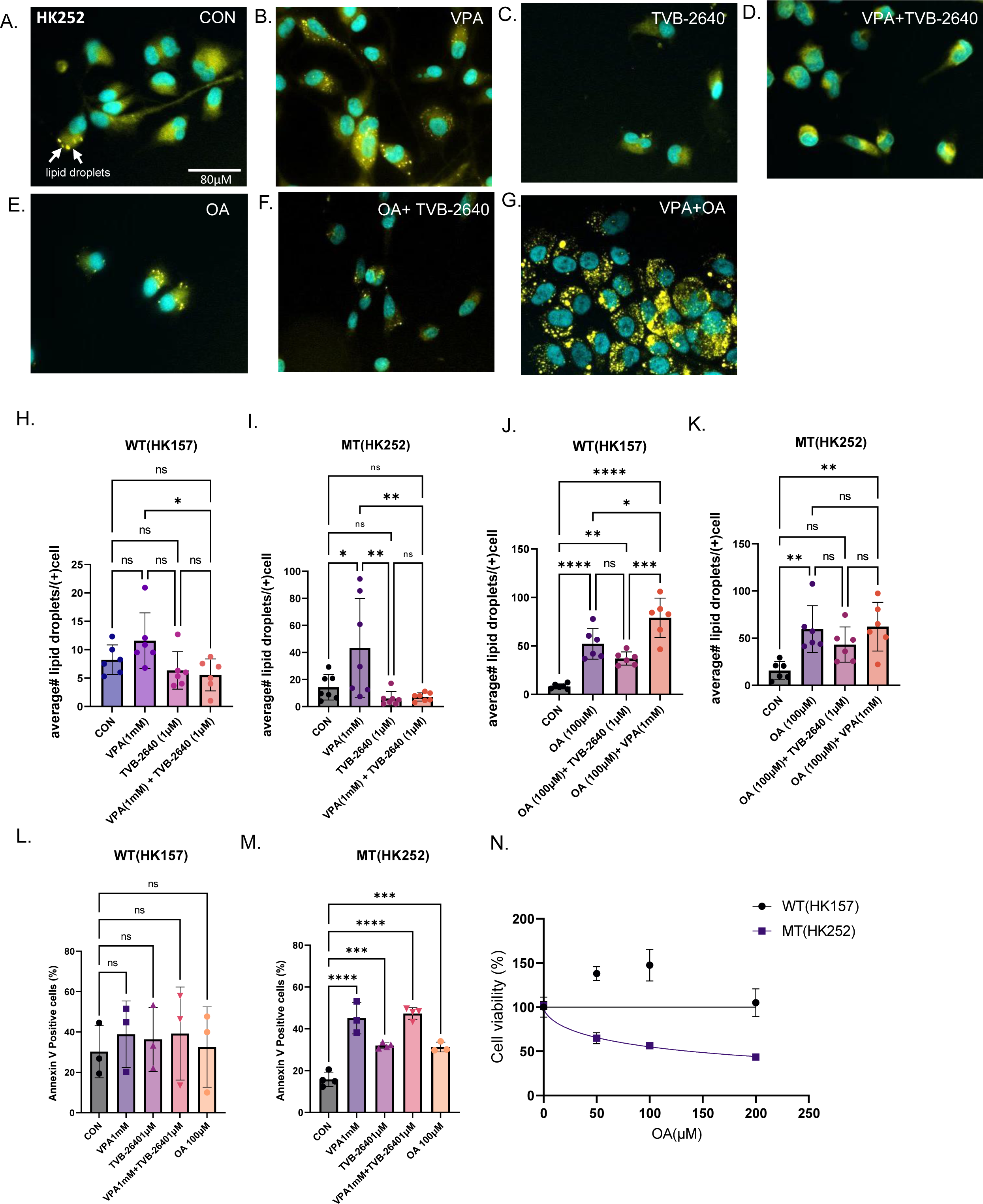
VPA and TVB-2640 alters lipid droplet formation and promotes apoptosis. A-G. Representative immunofluorescence images of lipid droplets in HK252 stained with Lipidspot^TM^ H-K. Quantification of lipid droplets per cell in each condition L-M. Flow cytometry quantification of Annexin V positive cells in HK252 and HK157 N. Cell viability of HK157 and HK252 after being treated with different concentration of oleic acid for 1 week. Non-Linear Regression. 2-way ANOVA, post hoc t-test, ****P value<0.0001; ***P value<0.001; **P value<0.01; **P value<0.05; Error bars ±SD

To determine whether lipid droplet formation is a protective mechanism or contributes to toxicity, we quantified the amount of annexin V positive cells after VPA, TVB-2640, and oleic acid treatment and found that, compared to control, all three treatments increased the proportion of annexin V positive cells only in the IDH1 MT cell line HK252 (Fig7L) but not in WT(Fig6M) cell line HK157. Oleic acid treatment also inhibited growth of IDH1 MT cell line HK252 but not WT cell line HK157 in a dose dependent manner (Fig7N). We also examined what fraction of cells that were positive for lipid droplets were also positive for annexin V by flow cytometry (SFig6A). In MT cell line HK252 we found that after VPA treatment about 30% of the annexin V positive cells were also positive for lipid droplets whereas 70% of annexin V positive cells were not positive for lipid droplets (SFig6B) suggesting that after VPA treatment some but not all cells are able to resist to VPA induced lipotoxicity by forming lipid droplets. Interestingly, TVB-2640 prevented lipid droplet formation in combination with VPA suggesting there might be some benefit of combining VPA with TVB-2640.

### HDACs are involved in regulation of lipogenic gene expression

Next, we wondered whether the effect of VPA on lipogenic genes is dependent or independent of HDAC inhibition. We re-analyzed RNA seq data from our recently published work ^15^ in which shRNAs used to knockdown different HDACs in IDH1 MT cell line HK252 to assess the effects on fatty acid metabolism genes. We found that knockdown of HDAC2,3,4, & 9 increased expression of lipogenic genes such FASN, stearyl coA desaturase (SCD) and HMGCR. HDAC1 knockdown slightly decreased expression of a few lipogenic enzymes but interestingly, only HDAC6 knockdown inhibited transcription of FASN. HDAC6 knockdown also inhibited transcription of other lipogenic enzymes such as ACACA, SCD and HMGCR but seemed to have a greater inhibitory effect on FASN (SFig7A). Using 2 different CRISPR sgRNAs, we prospectively knocked down HDAC6 in the HK252 cell line. HDAC6 is a microtubule deacetylase and as expected HDAC6 knockdown increased tubulin acetylation (SFig7B). Similarly, treatment of HK252 with VPA increased tubulin acetylation (SFig7C) in a dose dependent manner suggesting that VPA may mediate some of its effect on lipogenic genes through HDAC6. HDAC6 knockdown also inhibited transcription of FASN (SFig7D) and inhibited growth of MT cell line HK252 but not WT cell line HK157 (SFig7E). Together, we conclude the effect on lipogenic genes by VPA may be mediated through HDACs especially HDAC6. However, the upregulation of some of the lipogenic enzymes after HDAC knockdown potentially suggests a mechanism by which tumor cells might compensate and why some HDAC knockdowns inhibit growth of IDH1 MT glioma cells and others do not.

### FASN knockdown inhibits IDH1 mutant tumor growth in vivo and may enhance the effects of VPA treatment

Because tumor cells may be able to compensate for the loss of FASN function in vivo through the utilization of exogenous fatty acids, it was critical for us to determine the effect of diminished FASN function in vivo. It is currently unclear whether TVB-2640 can cross the blood brain barrier and whether there are any off-target effects of the drug. Hence, to test the effect of FASN inhibition in vivo we knocked down FASN in IDH1 MT cell lines BT142 and HK252 (Fig8A) using shRNAs. In vitro, FASN knockdown inhibited the growth of both HK252 (Fig8B) and BT142(Fig8C). In BT142, palmitate rescued some of the growth inhibitory effect of FASN knockdown with FASN shrna1 (Fig8D). We further found that FASN knockdown in combination with VPA was better at inhibiting growth of BT142 when compared to VPA and FASN knockdown alone (Fig8D) in vitro. Interestingly, palmitate did not rescue the growth inhibitory of FASN knockdown in combination with VPA. Therefore, we further hypothesized that in vivo combination of VPA with FASN knockdown may have a better growth inhibitory effect.

**Fig8:**
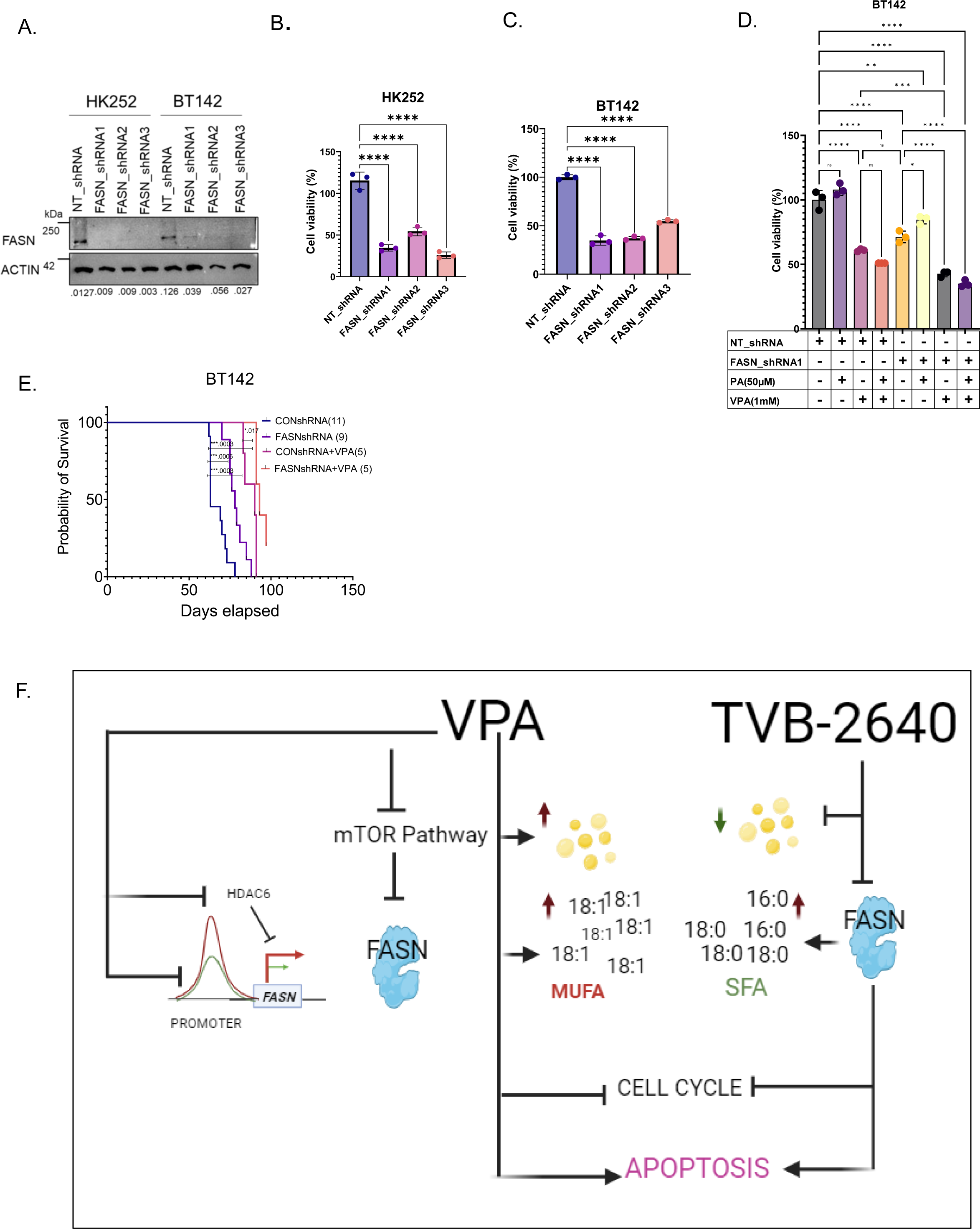
FASN knockdown enhances effect of VPA in vivo. A. Representative western blot showing FASN protein expression in HK252 and BT142 after FASN knockdown with three separate shRNAs. B-C. Relative cell viability of MT cell lines HK252 (B) and BT142 (C) after FASN knockdown. One-Way Anova, ****Pvalue <0.0001. D. Relative growth of BT142 with ctrl non target shRNA and FASN shRNA treated with VPA and PA for 1 week. E. Kaplan-Meier survival curve for mice treated with either saline or VPA (300mg/kg) implanted with NT BT142 or FASNshRNA1 KD BT142. ***P value<0.001; *P value<0.05 F. Schematic of mechanism of VPA and TVB-2640 in IDH MT glioma (created with bio render)

We found in vivo, compared to control, both VPA and FASN knockdown alone improved survival of mice (Fig8E) in the BT142 model. Further, the combination of VPA with FASN knockdown slightly, but significantly improved survival of mice when compared to VPA or FASN knockdown alone (Fig8E). These findings indicate that in vivo, even in the presumed presence of exogenous palmitate FASN knockdown alone can improve survival of mice. However, cancer cells can often use multiple ways to metabolically compensate or resist treatment hence knockdown of FASN may improve response to VPA in vivo.

## Discussion

In recent years several studies have emerged indicating that IDH1 MT gliomas are inhibited by HDACis such as panobinostat^15,21^ and belinostat^22^. In our study, we found that IDH1 MT gliomas are more sensitive to VPA when compared to IDH1 WT glioma cell lines. In addition to in vitro studies, we show that VPA improved survival of mice in vivo in two separate models of IDH1 MT glioma.

VPA is a drug with pleiotropic targets and, consistent with its role as an HDACi, VPA increased histone acetylation and transcriptionally activated many more genes compared to downregulated genes in both IDH1 WT and MT glioma cell lines. Surprisingly, we found that although VPA increased transcription, instead of opening up the chromatin, it led to diminished chromatin accessibility, suggestive of increased chromatin condensation. Interestingly, a recent study using human primary cells showed VPA promoted both histone acetylation and deacetylation, and the increase in histone acetylation with VPA is regional^23^. The increase in gene expression with VPA as we observed in our glioma cell lines may be driven by alternate promoters^24^ or complex interaction between promoter and enhancers^25^.

We identified several metabolic genes, many of which are involved in the regulation of de novo lipogenesis and cholesterol metabolism to be downregulated by VPA treatment. ATAC seq data further showed that many of these metabolic genes lost chromatin accessibility at the promoters after VPA treatment. Although VPA decreased chromatin accessibility in both WT and MT cell lines, the loss of promoter chromatin accessibility for lipogenic genes was significant in IDH1 MT but not IDH1 WT glioma cell lines. Previously, the disruption of fatty acid oxidation geneshas been found to decrease overall chromatin accessibility in the liver^26^; whether disruption of genes involved in lipogenesis also results in loss of chromatin accessibility is not known but we speculate that the disruption of lipogenic genes by VPA may contribute to the overall loss of chromatin accessibility seen in the IDH1 MT glioma cell lines.

We also found in both in IDH1 MT glioma cell lines and murine glioma model VPA inhibited the mTOR pathway, inhibited phosphorylation of S6 and downregulated FASN mRNA and protein expression. The effect of VPA on the mTOR pathway seemed to be less prominent on the WT cell lines in both models. Previously, VPA has been shown to inhibit phosphorylation of mTOR, AKT, S6 and promote autophagy in glioma^27^, gastric^28^ and prostate cancer cell lines^29^. We found treatment with rapamycin also inhibited FASN mRNA and protein expression in IDH1 MT GBM cell line. This is consistent with previous finding that showed that rapamycin targets lipid metabolism genes such as FASN^30^ and underscores that, at least in high grade IDH1 mutant tumors, the mTOR pathway plays a significant role.

Our data showed that both the FASN inhibitor TVB-2640 and VPA downregulates a suite of cell cycle and anti-apoptotic genes. FASN inhibition has previously been found to induce both cell cycle arrest^31^ and apoptosis^32^ in tumor cells. Our RNA seq data suggested inhibition of cell cycle processes with VPA treatment, and we also saw an increase in annexin V positive cells with VPA treatment suggesting VPA may exert its effect on both cell cycle and apoptosis via FASN downregulation. However, we cannot explain all the effects of VPA via FASN downregulation as VPA also targeted other lipogenic enzymes such as SCD and HMGCR. Our data suggested that inhibition of HMGCR also inhibited growth of IDH1 MT glioma cell lines, so it is possible that both inhibition of de novo lipogenesis and cholesterol metabolism contribute to the growth inhibitory effect by VPA.

Our lipidomic study suggested that although VPA targets FASN in IDH1 MT GBM cell line it doesn’t deplete palmitate. In fact, direct inhibition of FASN by TVB-2640 significantly increased free palmitic acid in IDH1 MT cell line HK252. In breast cancer cells, FASN inhibition has been found to increase unsaturated fatty acids, ceramides, and diacylglycerols^33^. In prostate cancer cells, inhibition of FASN by multiple FASN inhibitors increased synthesis of long chain unsaturated fatty acids and phospholipids^34^. This suggests that cancer cells potentially upregulate multiple other metabolic pathways to compensate for FASN inhibition and this may be conserved among multiple cancer types. Unlike TVB-2640, VPA inhibited multiple metabolic enzymes which may be the reason why we didn’t observe a dramatic increase in palmitic acid with VPA treatment. Interestingly, In IDH1 WT cell line HK157 we saw a decrease in palmitic acid with both VPA and TVB-2640, treatment suggesting that VPA may also work through FASN in some IDH1 WT cell lines. However, it is important to note that although direct inhibition of FASN inhibits growth of both WT and MT cell lines the downstream lipidome rewiring in WT and MT cell line is quite distinct.

The enzyme SCD is responsible for making oleic acid. Interestingly, we saw an increase in free oleic acid with VPA treatment in IDH1 MT cell line HK252 although VPA transcriptionally inhibited SCD. The increase in oleic acid could potentially be a compensatory mechanism but even if it is, it is most likely a failed compensatory mechanism, as our data suggested that treatment with oleic acid induced apoptosis and inhibited growth of IDH1 MT glioma cell line. This is further supported by a recent paper that showed that oleic acid shifted the balance towards monounsaturated fatty acids, causing apoptosis in IDH1 MT gliomas^35^. Although the prior work did not investigate lipid droplets, we found an increase in lipid droplets in IDH1 MT glioma cell line with both VPA and oleic acid treatment. Cancer cells generally form lipid droplets to prevent ER stress and lipotoxicity^36^. Hence, we cannot rule out the possibility that some IDH1 MT cells may escape treatment by inducing lipid droplet formation.

We also find that HDACs are not necessarily decoupled from the regulation of lipogenic enzymes. In fact, knockdown of some HDACs in IDH1 MT GBM cell line HK252 upregulated many lipogenic enzymes such as SCD and HMGCR. Our analysis showed a change in FASN mRNA expression with only HDAC6 knockdown in IDH1 MT glioma cell line HK252. VPA targets several HDACs and the fact that some HDAC knockdowns can upregulate metabolic genes is cautionary, as cancer cells can escape treatment through various mechanisms. It is possible that effects of VPA on other HDACs influence lipid metabolism in different ways, contributing to the growth inhibitory or compensatory effects. For example, two prior studies identified HDAC3 to be responsible for deacetylation of FASN^10,37^. HDAC3 knockdown in the liver was also found to reroute precursors towards lipid synthesis and sequestration into lipid droplets^38^.

Altogether, our study shows VPA reprograms both the chromatin and the lipidome of IDH1 MT glioma. Our in-depth analysis suggests VPA has many targets and may ultimately work through multiple mechanisms to inhibit growth of IDH1 MT gliomas and at least some of the mechanistic effect is mediated by FASN as summarized in Fig8F. We find some of the mechanistic actions of VPA on both IDH1 WT and IDH1 MT are the same while others are different. VPA increased transcription and led to chromatin condensation in both IDH1WT and IDH1 MT cell lines. However, we find the downstream effect on the lipidome is what distinguished IDH1 MT from IDH1 WT. Both VPA and the FASN inhibitor TVB-2640 inhibited growth of both IDH1 WT cell line HK157 and IDH1 MT cell line HK252, but both drugs induced apoptosis only in the IDH1 MT cell line but not in the IDH1 WT cell line. We also find that only IDH1 MT and not IDH1 WT is sensitive to oleic acid treatment and while VPA increased free oleic acid in IDH1 MT it decreased saturated free fatty acid in IDH1 WT cell line. Lastly, we also find VPA may mediate some of its effect on lipogenic enzymes via HDAC6. Interestingly, HDAC6 knockdown inhibited the growth of only IDH1 MT cell line HK252 but not IDH1 WT cell line HK157. We conclude VPA’s unique effect on the IDH1 MT lipidome may ultimately make them more sensitive to VPA than IDH1 WT.

Lipogenic enzymes can be significant therapeutic targets in cancer, but it is important to note that cancer cells can use multiple metabolic pathways for survival. In our study for instance, we found that only selective inhibition of FASN by TVB-2640 upregulated cholesterol metabolism in IDH1 MT gliomas. VPA, on the other hand, targeted enzymes that are important for both lipogenesis and cholesterol metabolism. Perhaps VPA’s pleotropic targets are more of an advantage than disadvantage for the treatment of IDH1 MT gliomas. Nonetheless, we found that VPA increased lipid droplet formation which might be a resistance mechanism and interestingly combination of TVB-2640 with VPA inhibited lipid droplet formation. In vivo, the combination of VPA with FASN knockdown was also better than VPA and FASN knockdown alone in improving survival of mice. Hence, we think in order to overcome the metabolic flexibility of cancer cells dual targeting of FASN by both VPA and TVB-2640 is potentially an effective therapeutic approach for IDH1 MT glioma.

## Methods

All experiments in this study comply with relevant ethical regulations and have been approved by the UCLA Institutional Review Board. Protocols for animal studies were approved by the institutional animal care and use committee at UCLA. All patient samples were de-identified and collected under informed consent and with the approval of UCLA Medical Institutional Review Board.

### Experimental cell lines and drugs

The primary patient derived IDH1 WT and IDH1 MT cell lines were established in our laboratory and cultured in serum free condition as previously described^39^. BT142 was purchased from ATCC. The murine glioma cell lines (NRAS G12V-shp53-shATRX-IDH11 wildtype; NPAC54B) and (NRAS G12V-R132H-shp53-shATRX-IDH11 mutant; NPAIC1) have been previously described (REF) and were cultured in the same media as patient-derived gliomaspheres. The cultures were regularly checked for mycoplasma and by STR to validate the source. The following drugs were used for the experiments. Valproic acid sodium salt,98% (Sigma-Aldrich, P4543-10G), panobinostat (Cayman, 13280), Belinostat (PXD101) (Sellechckem S1085), TVB-2640 (Sellechckem S9718), Oleic Acid (Cayman Chemical, 90260), Palmitic Acid (Sigma, 57-10-3), Rapamycin (LC laboratories, R5000).

### Animal Studies

Studies did not discriminate sex, and both male and females were used. Strains: 8 to 12-week-old NOD-SCID gamma null (NSG) mice (NOD.Cg-*Prkdc^scid^ Il2rg^tm1Wjl^*/SzJ Jackson Laboratory, 00557) were used to generate tumors from a patient-derived glioma line BT142. C57BL6 (Jackson Laboratory, 000664) were used to transplant a murine NPAC54B and NPAIC1 as described previously^40^. Firefly-luciferase-GFP BT142 (1X105) tumor cells were injected intracranially into the neostriatum of mice. 1 week after tumor cell line implant mice were treated with VPA (300mg/kg) of body weight twice a day every day of the week. For FASN knockdown and VPA experiment mice were treated with VPA (300mg/kg) of body weight twice a day 5 days a week. Treatment continued until the animal reached the endpoint. Tumor growth was monitored once a week by measuring luciferase activity using IVIS Lumina II bioluminescence imaging. Survival: Mice were euthanized if they looked hunched, lost significant body weight, or showed fur changes and decreased activity than normal for survival analysis.

### Bulk RNA sequencing and analysis

Qiagen RNeasy microkit was used for RNA extraction from gliomasphere. RNA quality was assessed using Bioanalyzer and only samples with a RIN score >8.0 were sequenced. RNA samples were pooled and barcoded, and libraries were prepared using TruSeq Stranded RNA (100 ng) + Ribozero Gold. Paired-end 2X75bp reads were aligned to the human reference genome (GRCh38.p3) using the STAR spliced read aligner (v 2.3.0e). Total counts of read fragments aligned to known gene regions within the human hg38 refSeq reference annotation was used as the basis for the quantification of gene expression. Differentially expressed genes were identified using EdgeR Bioconductor R-package, which are then considered and ranked based on False Discovery Rate (FDR Bejamini Hochberg adjusted p-values of ≤ 0.01(not 0.1?)). Gene Set Enrichment Analysis (GSEA) was carried out to determine the gene signatures differentially regulated in control and radiated cells and represented as heatmaps. R-Package V.3.2.5 (The R project for Statistical Computing, https://www.r-project.org/) was used to generate the PCA plots and heatmaps.

### ATAC sequencing and analysis

ATAC seq data was used from Garrett et al^15^. Alignment of reads was carried out using the Burrows-Wheeler Aligner mem using hg19 assembly. Peak calling was performed using MACS2 (with parameter setting –nomodel –shift 75), and differential peak analysis using featureCount and DESeq2 (default setting). Motif analysis and peak annotation was done using HOMER and GO analysis using HOMER (http://homer.ucsd.edu/homer/) and EnrichR (https://maayanlab.cloud/Enrichr/). UCSC Genome Browser was used to determine whether open regions displayed H3K27Ac and conserved TF binding sites. Integrated Genome viewer (IGV) was used to represent the peaks/open regions.

### Free fatty acid panel GC-MS & Analysis

1X10^6 million cells per sample per condition in triplicate were submitted to UCSD Lipidomic Core. Samples were prepared using previously published protocol Quehenberger et al. (2010)^41^ and analyzed at UCSD Lipidomics Core.

### Cell viability assay

WT and MT cells were plated at a density of 5000 cells per well in 96-well plates. Proliferation was assessed 7 days after treatment with respective drugs using CellTiter-Glo® luminescent Cell Viability Assay (Fisher Scientific, PRG9242). The luminescence signal was measured in a luminometer, on day 0 and day7. Luminescence signal was normalized to initial reading at day0.

### Western blot assay

At experimental end points cells were collected and lysed in RIPA buffer with protease and phosphatase inhibitors. Cells were kept on ice for 10 mins and centrifuged at full speed for 5 minutes. Protein concentration was measured by Bradford assay using a BSA standard. Protein lysates were run on a SDS-PAGE on 4%–12% gradient polyacrylamide gel (Thermo-fischer Scientific) and transferred onto nitrocellulose membranes. Blots were incubated with primary antibody diluted in 5% milk overnight. Blots were washed and incubated with HRP-conjugated secondary antibodies. Quantitation of protein levels was performed in ImageJ by normalizing to loading control, β-actin. Antibodies FASN (CST, 3180), PS6235/236 (CST, 2211), S6 (CST, 2217), BACTIN (CST, 4970), HDAC6 (CST, 7558), H3K27AC (CST, 8173), H3 (CST 9717).

### Annexin V/7ADD assay

Cells were treated for 4 days with VPA and TVB-2640 and cells were prepared according to manufacturer’s instruction for Biotium Annexin V and 7-AAD Apoptosis Kit and acquired by flow cytometry within an hour.

### Intracellular D-2-HG Quantification

Intracellular D-2-HG was quantified using enzymatic assays originally described by Balss, *et al.* 2012, and who we thank for generously providing key reagents that are also commercially available in the D-2-HG Assay Kit (*MilliporeSigma*, Catalog #MAK320, Burlington, MA, USA). All experiments were performed in triplicate, unless otherwise specified. Briefly, cells were harvested and lysed prior to splitting each sample into two aliquots for D-2-HG quantification and total protein quantification using the Pierce BCA Protein Assay Kit (*Thermo Fisher Scientific*, Catalog #23225, Waltham, MA, USA), respectively. Deproteination of the D-2-HG quantification aliquot was achieved using 3.0 mL of Proteinase K (*Qiagen*, Catalog #19131, Hildren, Germany) per 100.0 mL of cell lysis solution. 25.0 mL of lysate was then added to 75.0 mL of assay solution containing 0.1 mg of the enzyme D-2-HG dehydrogenase (HGDH), 100.0 mM NAD+, 5.0 mM resazurin, and 0.01 U/mL diaphorase in 100.0 mM HEPES (pH 8.0). In the presence of D-2-HG, HGDH converts D-2-HG to α-ketoglutarate (α-KG) in a NAD+ dependent manner. The reduction of NAD+ to NADH enables the conversion of resazurin to fluorescent resorufin, which was then quantified via fluorometric detection (λ_ex_ = 540 nm, λ_em_ = 590 nm) on a Wallace Victor2 1420 Miltilabel HTS Counter (*PerkinElmer*, Waltham, MA, USA). D-2-HG quantification (pmole/mg protein) of a given sample was based on a standard curve of known D-2-HG concentrations. Two-tailed Student’s *t*-tests were used to compare means between groups, with statistical significance set at *P<*0.05.

### Lipid droplet staining and flow cytometry

LipidSpot Lipid Droplet Stain 647 was used to stain lipid droplets in gliomasphere cultures. Cells were plated on a 24 well dish in Cultrex UltiMatrix covered coverslips. Cells were treated with drugs for 4 days and at experimental endpoint, media was removed, cells were washed, and incubated with 1X dye diluted in DPBS for 15 min at 37° C. After 15 mins the dye was removed, and cells were fixed with 4% PFA. Nuclei were stained with Hoechst and mounted on coverslips with vector shield. Coverslips were imaged and analyzed using Image J.

For flow cytometry quantification at experimental endpoint media was removed and cells were incubated with 1X dye diluted in DPBS for 30 min at 37° C. After 30 min cells, the dye was removed, cells were dissociated into single cells and acquired using flow cytometry.

### shRNA and CRISPR/Cas9 Lentiviral Knockdown

The following plasmids were used for knockdown experiments Scramble shRNA plasmid # 1864 (Addgene); FASN shRNA1 plasmid # 82327(Addgene); FASN shRNA2 _ TRCN0000002125 (Horizon); FASN shRNA2 _ TRCN0000002126 (Horizon); FASN_shRNA3_TRCN0000002127 (Horizon); HDAC6 sgRNA 5488: CAAGGAGCAACTGATCCAGG, HDAC6 sgRNA 5501: CCTAGATCGCTGCGTGTCCT, AAVS1 sgRNA: GGGCCACTAGGGACAGGAT were cloned into pLentiCRISPRv2. lentiCRISPR v2 was a gift from Feng Zhang (Addgene plasmid # 52961; http://n2t.net/addgene:52961; RRID: Addgene_52961)^42^.

These plasmids were transfected into 293T cells along with a 2nd generation viral DR8.74 package and VSV-g envelope for production of lentivirus. Cells were infected with lentivirus and puro selected for a week before doing growth experiments. For cells infected with FASN shRNA media was supplemented with palmitic acid (50µM) along to puromycin to prevent all knockout cells from dying.

### Statistical Analysis

All data are expressed as mean ± SD. *P* values less than 0.05 were taken as significant and were calculated in Graph Pad Prism 9.0 using one-way ANOVA for multiple comparison with Bonferroni correction, followed by post-hoc t-test. Log-rank analysis was used to determine the statistical significance of Kaplan-Meier survival curves. R-package was used for statistical analysis of sequencing experiments. Schematics used in the figures were generated in Biorender.

### Data availability

All raw sequencing data will be available in the NCBI Gene Expression (prepared prior to publication).

## Acknowledgements

The authors thank the UCSD Lipidomic Core, JCCC Flow Cytometry Core, UNGC, and the TCGB core for technical contributions.

## Author contribution

L.S.E and H.I.K conceptualized the study. L.S.E, M.C, A.G.A, B.G, H.Q, T.L, M.C.G performed experiments. R.K, Y.Q, L.S.E performed bioinformatics analysis. A.L., M.G.C, P.R.L provided reagents and cell lines. L.S.E and H.I.K wrote and edited the manuscript. All authors contributed to the revision of the manuscript.

---

**SFig1.**
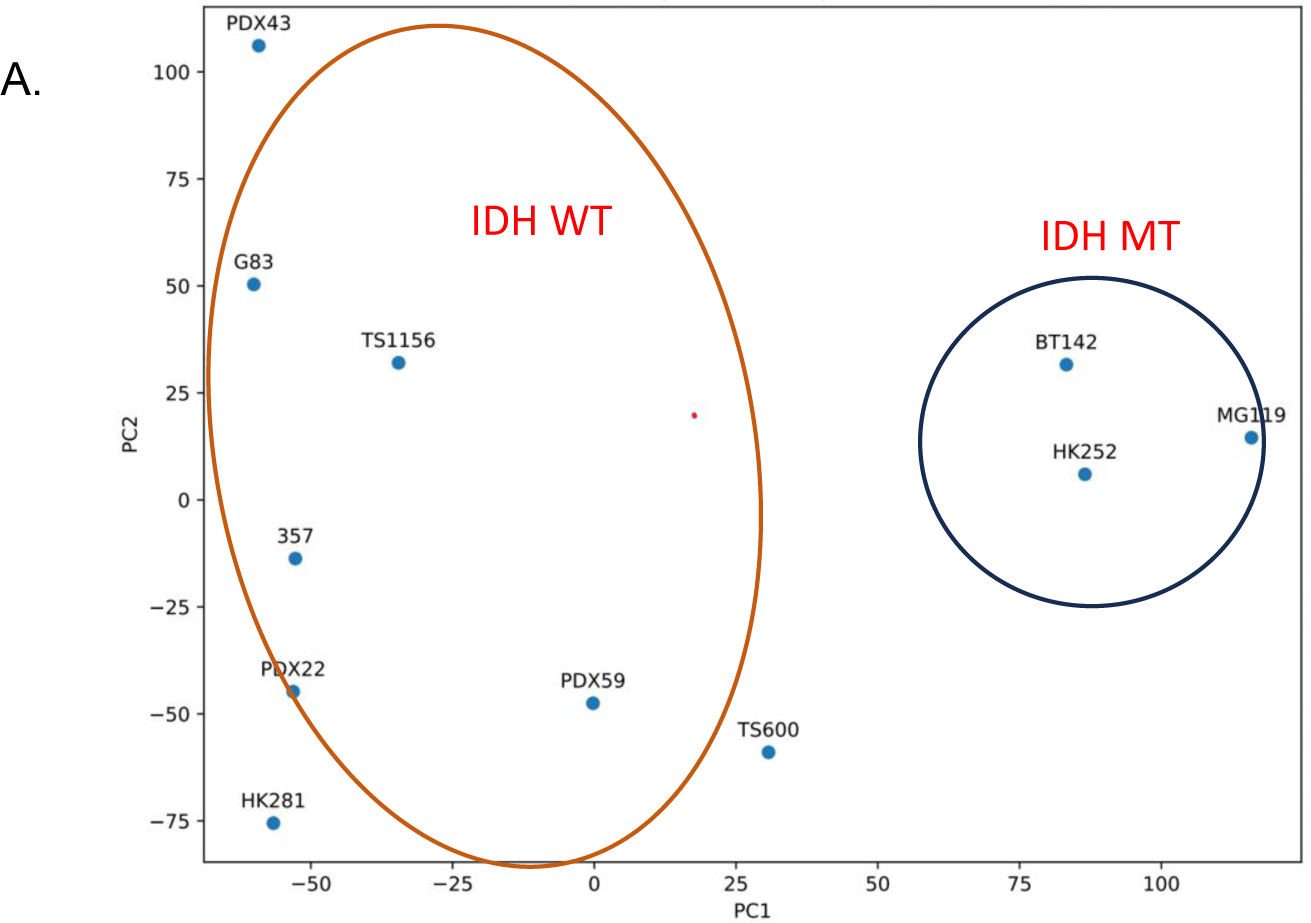
The transcriptome of BT142 is similar to heterozygous IDH MT cell line and distinct from IDH WT cell lines. A. Principal component analysis of bulk RNA sequencing of IDH MT and IDH WT cell lines. HK252, BT142 and MG119 are IDH MT cell lines and clusters together separately from all the IDH WT cell lines.

**SFig2.**
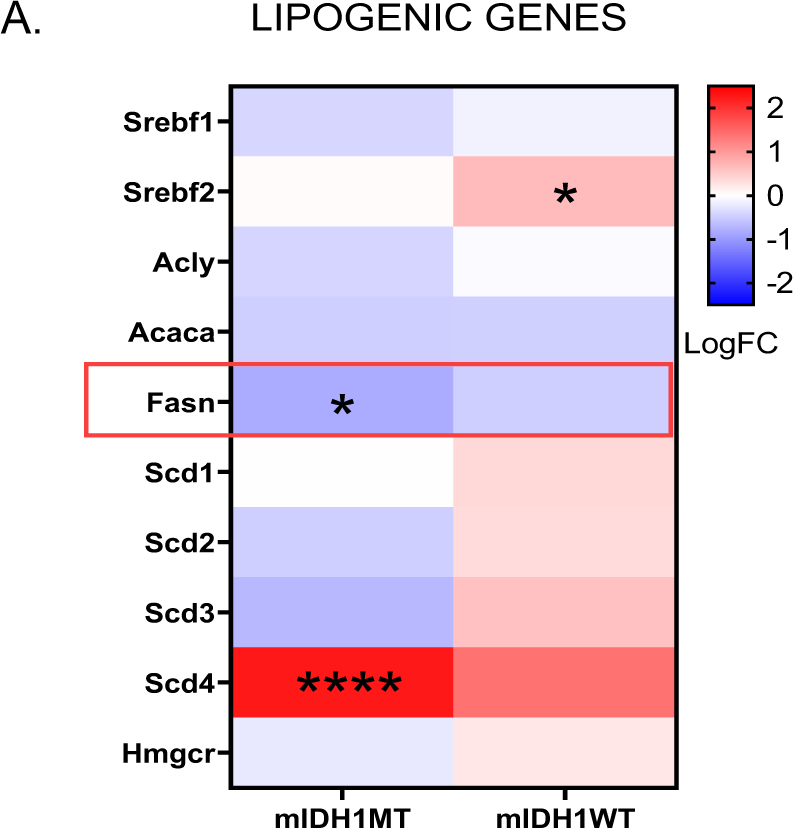
VPA alters lipogenic gene expression in murine isogenic model of IDH MT GBM. A. Selected lipogenic genes that are downregulated in mIDH1WT and mIDH1MT cell lines. ****P value<0.0001; P value<0.05;

**SFig3:**
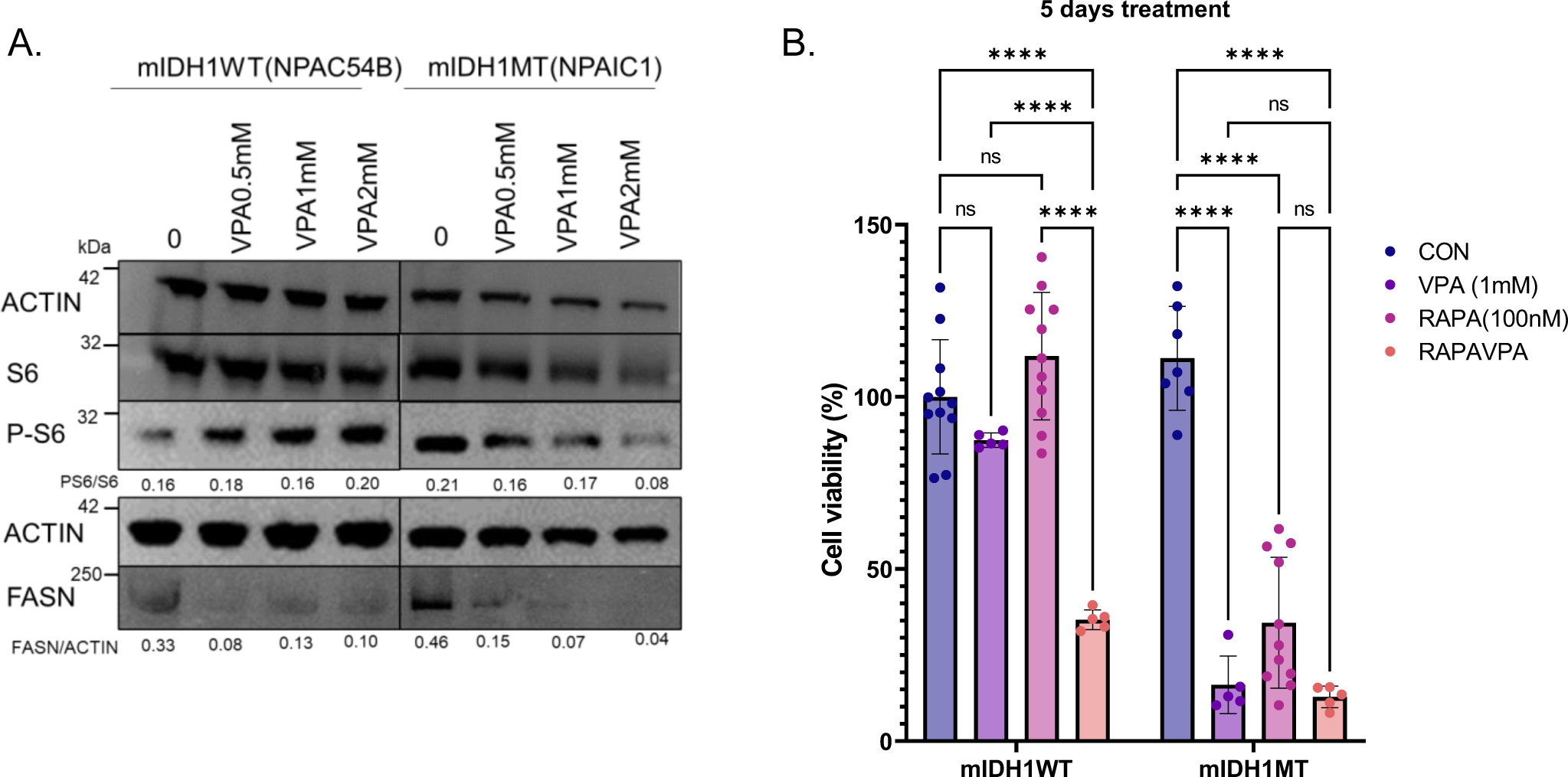
VPA decreases PS6 and FASN protein expression in mIDH1MT (NPAIC1) cell line. A. Representative western blot of PS6 and FASN in NPAC54B and NPAIC1 treated with increasing concentration of VPA for 4 days.

B. Relative cell viability after treatment with VPA, RAPA, & combination for 5 days. 2-way ANOVA, post hoc t-test,****P value<0.0001; Error bars ±SD.

**SFig4:**
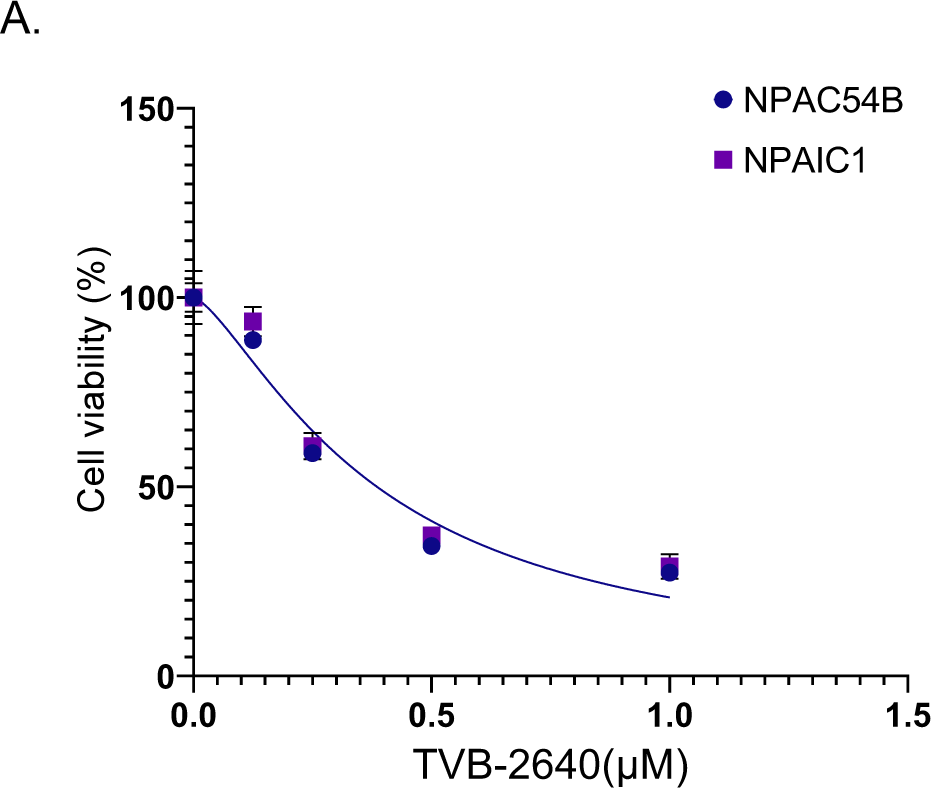
Both NPAIC1 and NPAC54B respond to TVB-2640. A.Dose response curve for NPAIC1 and NPAC54B treated with TVB-2640, Non Linear Regression.

**SFig5:**
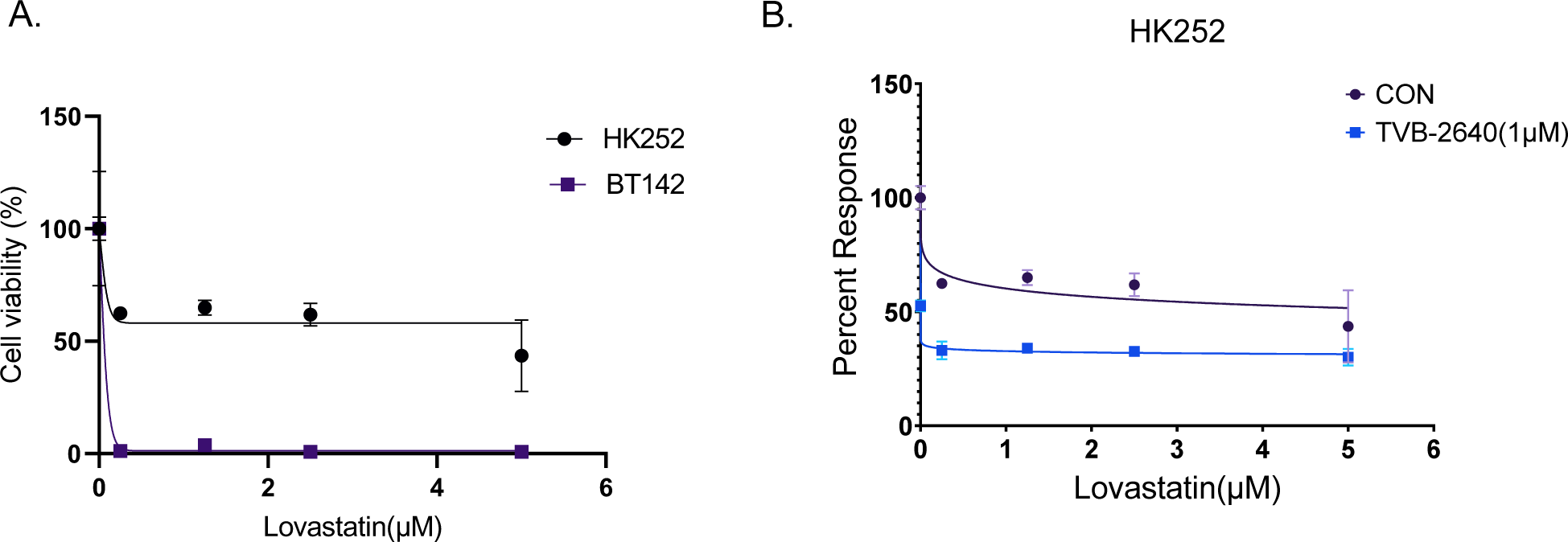
Lovastatin inhibits growth of IDH MT cell lines. A. Relative cell viability after treatment with lovastatin for 1 week. Non-Linear Regression.

**SFig6:**
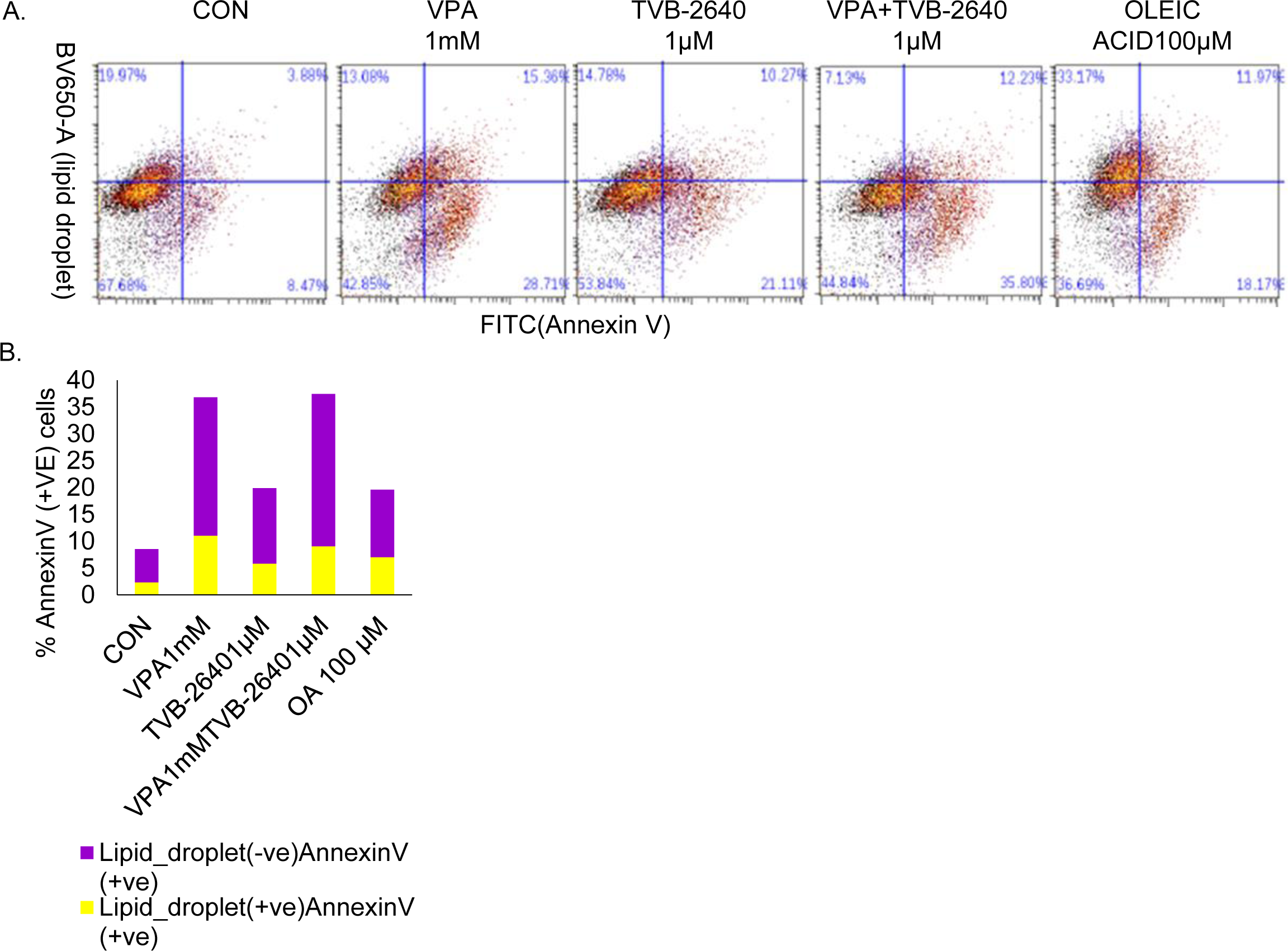
Relative proportion of cells that are positive for both lipid droplet and annexin V. A. Representative flow cytometry traces of cells stained with Annexin V and Lipid droplets. Cells were treated for 4 days. B. Quantification of proportion of annexin V and lipid droplet double positive cells.

**SFig7:**
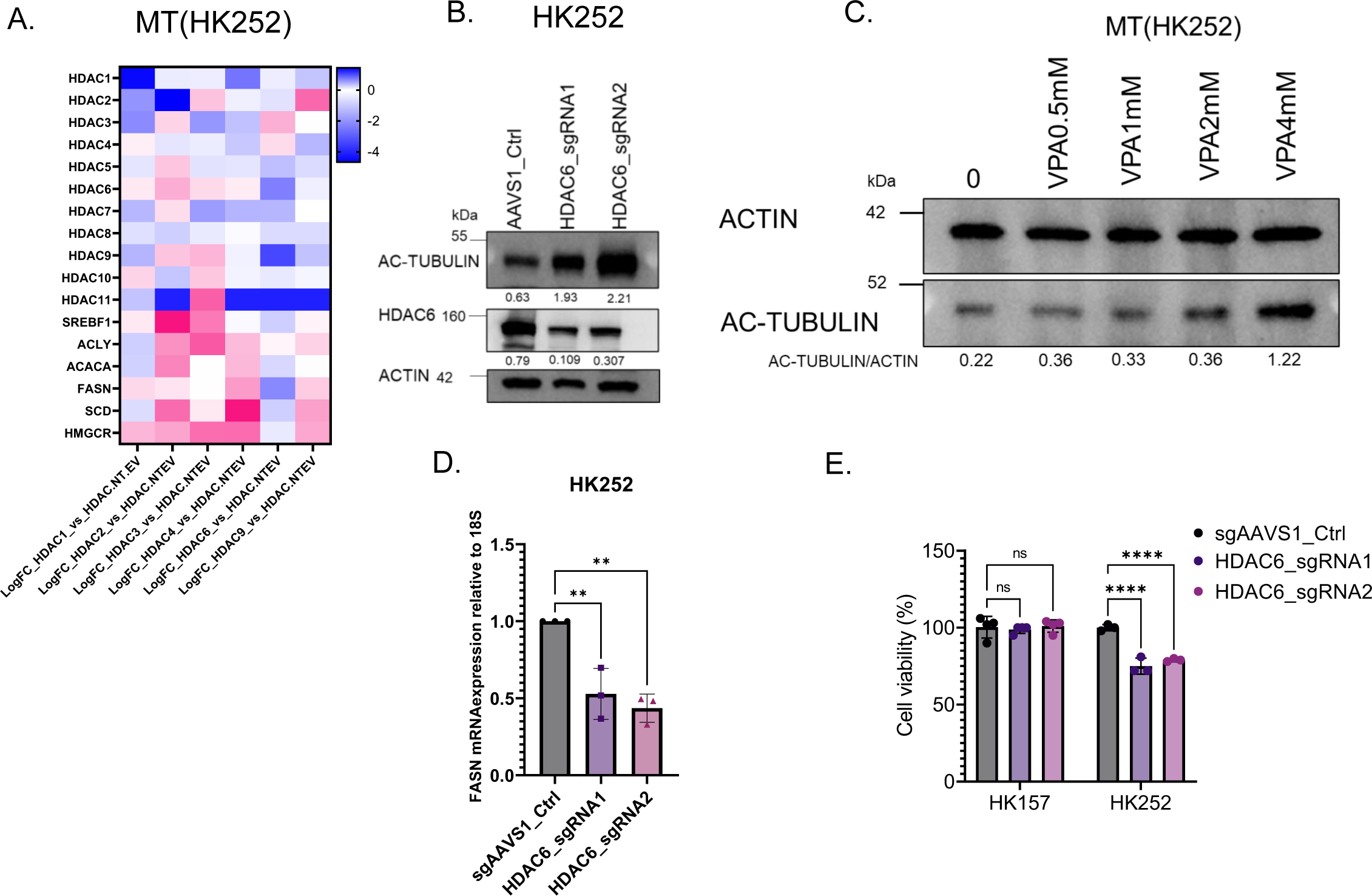
HDACs are involved in regulation of lipogenic genes in IDH MT GBM. A. Heatmap showing expression of lipogenic genes after HDAC knockdowns B. Representative WB showing tubulin acetylation and HDAC6 protein expression after HDAC6 knockdown with two CRISPR sgRNA C. Representative WB showing acetylation of tubulin after treatment with VPA for 4 days D. FASN mRNA expression after HDAC6 knockdown E. Relative cell viability of HK252 and HK157 after HDAC6 knockdown. 1-way ANOVA, **P value< 0.01, ****P value<0.0001;

